# Structural Requirements for Activity of Mind bomb1 in Notch Signaling

**DOI:** 10.1101/2024.03.01.582834

**Authors:** Ruili Cao, Oren Gozlan, Lena Tveriakhina, Haixia Zhou, Hanjie Jiang, Philip A. Cole, Jon C. Aster, David Sprinzak, Stephen C. Blacklow

## Abstract

Mind bomb 1 (MIB1) is a RING E3 ligase that ubiquitinates Notch ligands, a necessary step for induction of Notch signaling. The structural basis for binding of the JAG1 ligand by the N-terminal region of MIB1 is known, yet how the ankyrin (ANK) and RING domains of MIB1 cooperate to catalyze ubiquitin transfer from E2∼Ub to Notch ligands remains unclear. Here, we show that the third RING domain and adjacent coiled coil region of MIB1 (ccRING3) drives MIB1 dimerization and that ubiquitin transfer activity of MIB1 relies solely on RING3. We report x-ray crystal structures of a UbcH5B-ccRING3 complex as a fusion protein and of the ANK region. Directly tethering the N-terminal region to ccRING3 forms a minimal MIB1 protein, which is sufficient to induce a Notch response in receiver cells. Together, these studies define the functional elements of an E3 ligase needed for ligands to induce a Notch signaling response.

## Introduction

Notch signaling is a conserved system of cellular communication that plays a pivotal role in development and adult tissue homeostasis.^1–3^ Signals transduced by Notch receptors influence cell fate decisions in many tissues, and mutations of various core protein components of the Notch pathway result in developmental disorders of the gastrointestinal, cardiovascular, hematopoietic, and central nervous systems.^3^

Notch signaling is initiated when a transmembrane ligand on a signal-sending cell binds a Notch protein on a receiver cell. Bound ligand then induces proteolytic cleavage of Notch at a juxtamembrane extracellular site by an ADAM metalloprotease at site S2, which enables subsequent proteolytic processing of the truncated Notch protein at the inner membrane leaflet by gamma secretase^4,5^ at site S3 and release of the Notch intracellular domain (NICD) into the cell. Entry of NICD into the nucleus then leads to assembly of a transcriptional activation complex that induces the expression of Notch target genes.^6–9^

A crucial event that links the formation of ligand-receptor complexes to productive signaling is the requirement for ubiquitination-dependent endocytosis of the ligand in sender cells.^10^ Genetic investigations in flies and zebrafish identified two distinct E3 ubiquitin ligases, Mind bomb (MIB) and Neuralized (NEUR), that can catalyze the transfer of ubiquitin to the cytoplasmic tails of Notch ligands.^11–13^ MIB and NEUR have neither sequence nor structural similarity and they appear to recognize different sequences within the cytoplasmic tails of Notch ligands.^14^ The two E3 ligases, however, can functionally substitute for each other in certain cellular contexts,^15^ suggesting a degree of functional redundancy between them.

In mammals, there are two MIB proteins capable of ubiquitinating Notch ligands, MIB1 and MIB2. ^12,16^ They are modular proteins that contain an N-terminal substrate recognition domain encompassing MZM and REP regions,^17^ a central ankyrin repeat domain (ANK), and a C-terminal region that includes a series of RING domains (Figure 1A). The main difference between MIB1 and MIB2 is that MIB1 has three RING domains, whereas MIB2 has only two, with its second RING domain homologous to RING3 of MIB1.

**Fig. 1.**
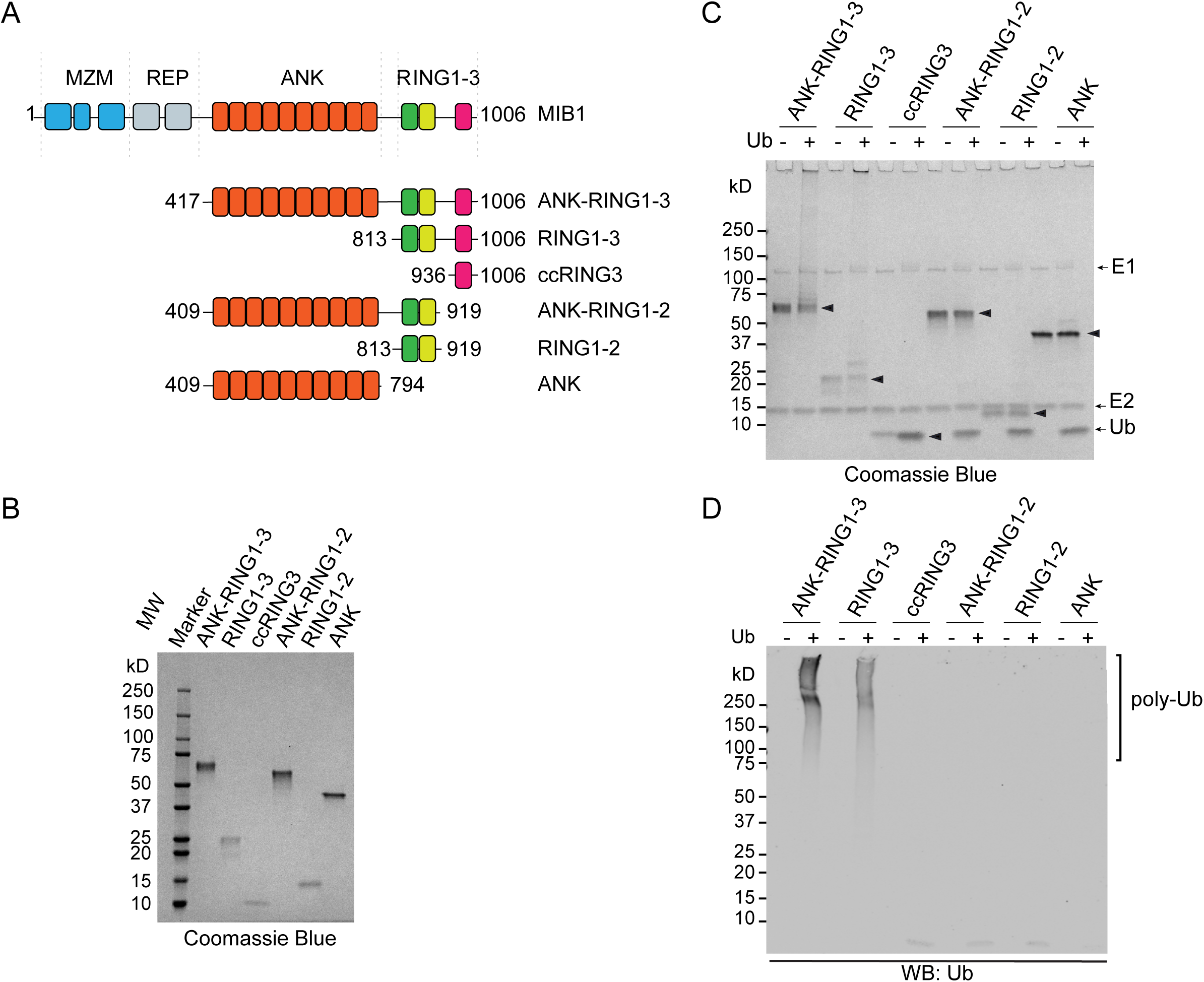
The ccRING3 region is responsible for the ubiquitination activity of MIB1. A, Domain organization of MIB1 and design of truncated MIB1 protein variants tested in the ubiquitination assay. B, SDS-PAGE gel of purified MIB1 truncation variants, visualized by staining with Coomassie Blue. C and D, SDS-PAGE and Westen Blot analysis of *in vitro* auto-ubiquitination activity of truncated MIB1 variants. Reactions were performed for 3h in the absence (-) or presence of Ub (+). Products were separated by SDS-PAGE, and detected either by Coomassie Blue staining (C), or with an anti-Ubiquitin (Ub) antibody (D). Arrowheads in panel C indicate the migration positions of the unmodified different MIB1 variants.

Functionally, MIB1 is essential for Notch signaling during mammalian development, whereas MIB2 appears to be dispensable. MIB2 knockout mice are viable and grossly normal, but MIB1 knockout mice die in utero at roughly embryonic day 10.5.^18–20^ Conditional inactivation of MIB1 in different tissues also results in Notch loss of function phenotypes, again highlighting its importance in developmental Notch signaling.^21–24^

MIB1 is also implicated in other signaling pathways and cellular processes. It influences Wnt/β- catenin signaling by ubiquitinating the receptor-like tyrosine kinase RYK, facilitating the endocytosis of RYK in response to Wnt signaling.^25^ It is also linked to centriolar satellite formation and ciliogenesis, adenovirus infection, NF-kB activation, and signaling events that lead to cell death.^26–30^ These other activities of MIB1 highlight its importance as a regulator of other cellular processes beyond Notch signaling.

Although previous work has elucidated the basis for substrate recognition by the N-terminal, MZM/REP region of MIB1,^17^ how the modular structural elements of MIB1 work together to coordinate the transfer of ubiquitin to its substrates is less clear. In the work reported here, we show that the ubiquitin transfer activity of MIB1 relies on RING3, but not on RING1 or RING2, and that formation of MIB1 dimers relies on RING3 and an adjacent coiled coil region, together termed ccRING3. We determine x-ray crystal structures of the ANK domain and of a complex of ccRING3 with UbcH5B, a functional E2 subunit for MIB1, as a fusion protein, and use negative staining electron microscopy to image full-length MIB1, identifying flexibility in the linkages between modular elements. Lastly, we investigate the structural elements of MIB1 required to stimulate Notch signals, and find that directly tethering the N-terminal, MZM/REP region to ccRING3 in a minimized MIB1 protein (mini-MIB1) is sufficient to induce a Notch signal in receiver cells. Together, these studies define the functional elements of an E3 ligase needed for ligands to induce a Notch signaling response.

## Results

### RING3 is responsible for the ubiquitination activity of MIB1

MIB1 contains three RING domains at its C-terminal end. We analyzed the autoubiquitination activity of a series of truncated MIB1 variants encompassing the ANK domain and the three RINGs (Fig. 1A) to determine which RING domain(s) are necessary for ubiquitin transfer activity. We purified each protein to homogeneity, as judged by SDS-PAGE (Fig. 1B), and performed an autoubiquitination assay for each protein in the presence of E1, the E2 UbcH5B, and ubiquitin (Fig. 1C&D; Supplementary Fig 1). Whereas ANK-RING1-3 and RING1-3 proteins showed robust self-ubiquitination activity in the presence of E1, E2, and Ub, the two proteins lacking ccRING3, labeled ANK-RING1-2 and RING1-2, did not self-ubiquitinate, indicating that ccRING3 is absolutely required for self-ubiquitination. In isolation, purified ccRING3 did not self-ubiquitinate, likely because there is not a good lysine acceptor within the isolated ccRING3 polypeptide.

### Structure of a UbcH5B-ccRING3 fusion protein

Efforts to determine the structure of an E2-ccRING3 complex using separately purified ccRING3 and UbcH5B proteins was not successful. Therefore, we constructed a fusion protein with UbcH5B at the N-terminus connected to the ccRING3 portion of MIB1 at residue 936 with an 18-residue glycine-serine (GS) linker (Fig 2A). After purification and crystallization of this fusion protein, we determined its x-ray crystal structure to 2.4 Å resolution (Table S1).

**Fig. 2.**
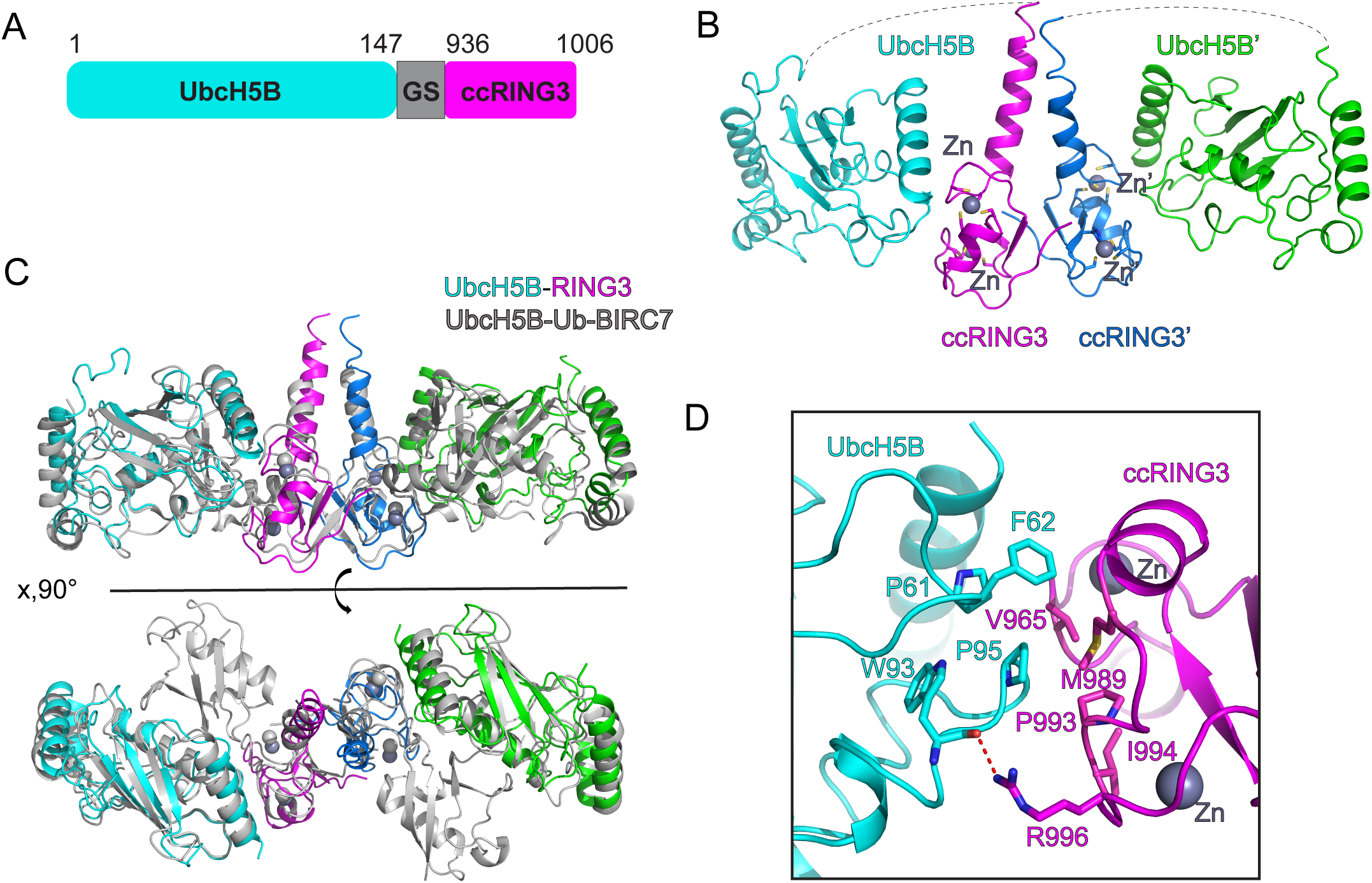
Structure of a UbcH5B-ccRING3 fusion protein. A, Schematic of the fusion protein used to mimic a UbcH5B-ccRING3 complex, in which UbcH5B was tethered to ccRING3 (residues 936-1006) using a Gly-Ser (GS) linker. B, Cartoon representation of the structure of the UbcH5B-ccRING3 dimer of heterodimers. The ccRING3 subunits are magenta and blue, and their partner UbcH5B subunits are cyan and green, respectively. Zn^++^ ions are colored gray and rendered as spheres. C, Comparison of the UbcH5B-ccRING3 dimer (colored as in B) with the UbcH5B-Ub-BIRC7 dimeric complex (gray). D, Contacts between ccRING3 of MIB1 and UbcH5B. Residues that approach within van der Waals distance are rendered as sticks, and a hydrogen bond from W93 of UbcH5B to the R996 side chain of MIB1 is shown as a dotted red line.

In the structure, two molecules of ccRING3 form a homodimer, with each copy bound to one UbcH5B molecule in a complex with 2:2 stoichiometry (Fig. 2B). The structure of each MIB1 subunit of the complex closely resembles that of BIRC7 in complex with UbcH5B-ubiquitin (Fig. 2C), as well as that of other dimeric RING E3 ligase domains complexed with E2 or E2-ubiquitin.^31–33^ Dimerization is mediated both by coiled-coil contacts and hydrophobic packing between the RING3 domains. Residues V965, M989, P993 and I994 of the RING3 domain, which are highly conserved among RING E3 ligases, form hydrophobic interactions with P61, F62 and P95 of UbcH5B, as seen in other complexes between E2 proteins and RING E3s (Fig. 2D, Supplementary Fig. 2). R996 of the RING3 domain also forms a hydrogen bond with the main chain of W93 of UbcH5B (Fig. 2D).

### Structure of the ANK repeat domain

We produced the ANK repeat domain (residues 409-794) in bacteria, purified it to homogeneity, grew diffracting crystals and determined its structure to 2.4 Å resolution using molecular replacement (Table S2). The model includes 10 ankyrin repeats, which adopt a horseshoe-like arrangement (Fig. 3A), extending 90 Å across its length from residue 428 at the N-terminus of the first repeat to position 794 at the C-terminal end. The ninth repeat features an atypically extended alpha helix, which participates in crystal packing interactions (Supplementary Fig. 3A). Strikingly, the concave face of the ANK domain is highly conserved, suggesting functional importance (Supplementary Fig. 3B).

**Fig 3.**
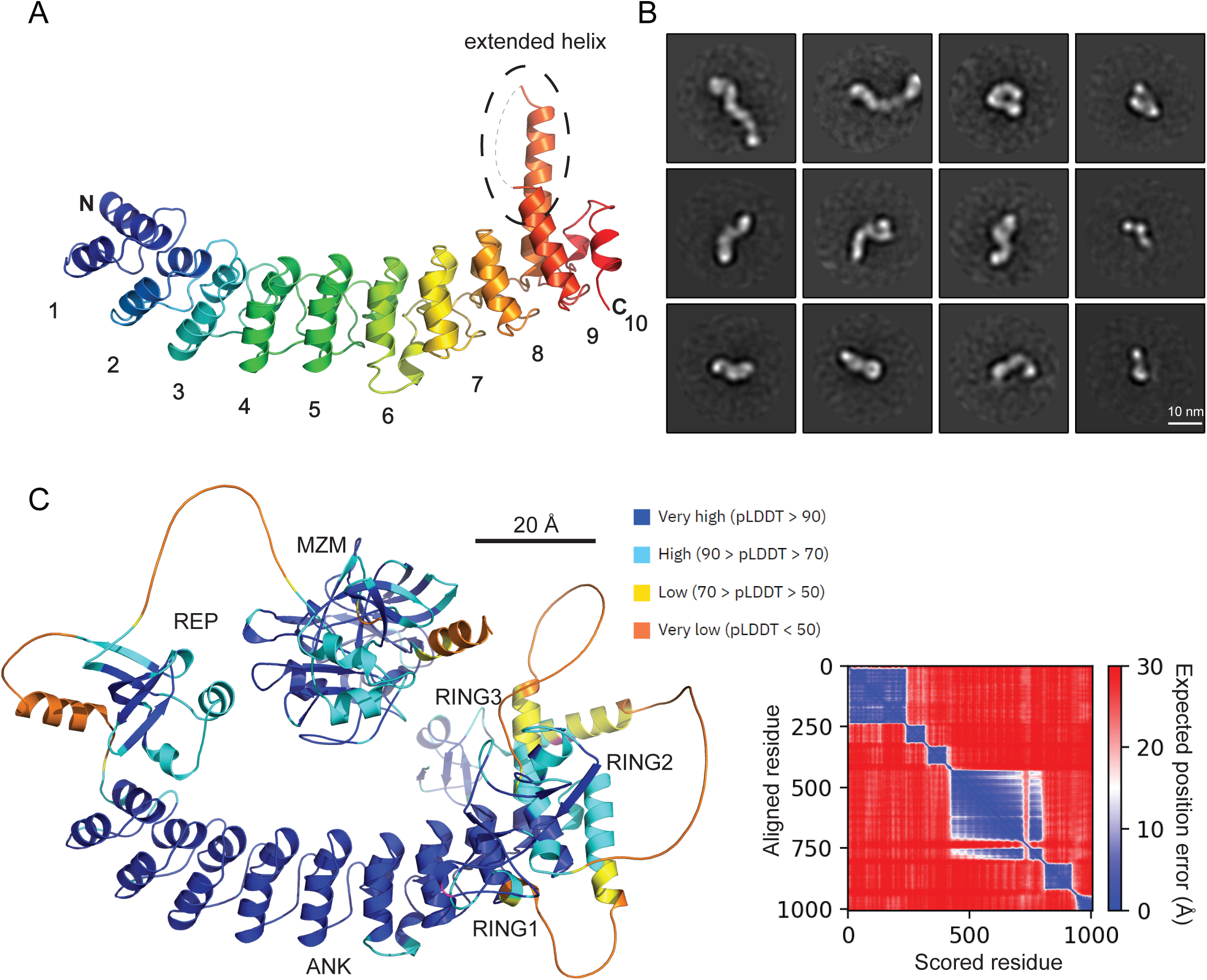
Structure of the ANK region and negative stain EM of full length MIB1. A, Cartoon representation of the MIB1 ANK domain structure, colored from blue (N-terminus) to red (C-terminus). B, 2D class averages of negative-stain images of full length murine MIB1, illustrating the heterogeneity of conformations of the full-length protein. Scale bar, 10 nm. C, Cartoon representation of the Colabfold^34^ model of MIB1 (left panel), colored by pLDDT value (per residue model confidence score) on a sliding scale from blue (pLDDT >90) to orange (pLDDT < 50). The right panel shows the expected position error plot, colored on a sliding scale from dark blue (0 Å) to red (>30 Å), for the Colabfold model of full-length MIB1, consistent with a lack of long-range interactions in the protein. Scale bar, 20 Å (2 nm).

### Negative stain electron microscopy of purified, full-length, murine MIB1 suggests interdomain flexibility

We purified full-length murine MIB1 and examined it using negative stain electron microscopy (EM) because the yield of full-length human MIB1, which we used for ubiquitination assays (see below), was insufficient for structural studies. The EM images show that MIB1 exhibits structural heterogeneity and adopts a variety of different conformations with extended and closed forms visible among the two-dimensional class averages (Fig. 3B), suggesting that MIB1 is highly dynamic and has flexible linkers connecting individually structured elements. The Alphafold2^34^ model for full-length MIB1 is consistent with this interpretation, with low expected position error (EPE) values for residue pairs within the structured domains and large EPE values for residue pairs located in different domains (Fig. 3C). The size of the particles also suggests that full-length MIB1 is dimeric like the isolated ccRING3 domain (see below).

### Mutational structure-function analysis of MIB1

We examined the consequences of mutations of MIB1 (Supplementary Fig. 4A) on the ability of Delta-like 4 (DLL4) to induce a transcriptional response. This activity was tested using a co-culture assay in which DLL4 expressing sender cells were co-cultured with U2OS receiver cells expressing a Notch1-gal4 chimeric protein and a UAS-regulated luciferase reporter gene^35^ (Fig. 4A). As sender cells we used U2OS MIB1^-/-^ knockout cells, stably transfected with DLL4 alone, wild-type (wt, Full-length) MIB1 alone, or with DLL4 and either wt or mutated MIB using lentiviral transduction. We confirmed that all proteins were expressed in amounts comparable to or greater than full-length wt MIB1 (Supplementary Fig. 4B). Addition of DLL4 to U2OS MIB1^-/-^ knockout cells did not induce a transcriptional response, confirming the requirement of MIB1 for the induction of a reporter signal (Fig. 4A). As expected, delivery of wt (Full-length) MIB1 and DLL4 into MIB1^-/-^ knockout sender cells leads to substantial reporter gene induction in Notch1-gal4 receiver cells, whereas delivery of MIB1 lacking the MZM/REP region (ΔMZM/REP), which is required for binding of ligand cytoplasmic tails,^17^ fails to rescue reporter activity. Likewise, deletion of the ccRING3 region also renders MIB1 unable to rescue signaling activity in the reporter assay (Fig. 4A), consistent with the requirement for ccRING3 in ubiquitin transfer (Fig. 1).

**Fig. 4.**
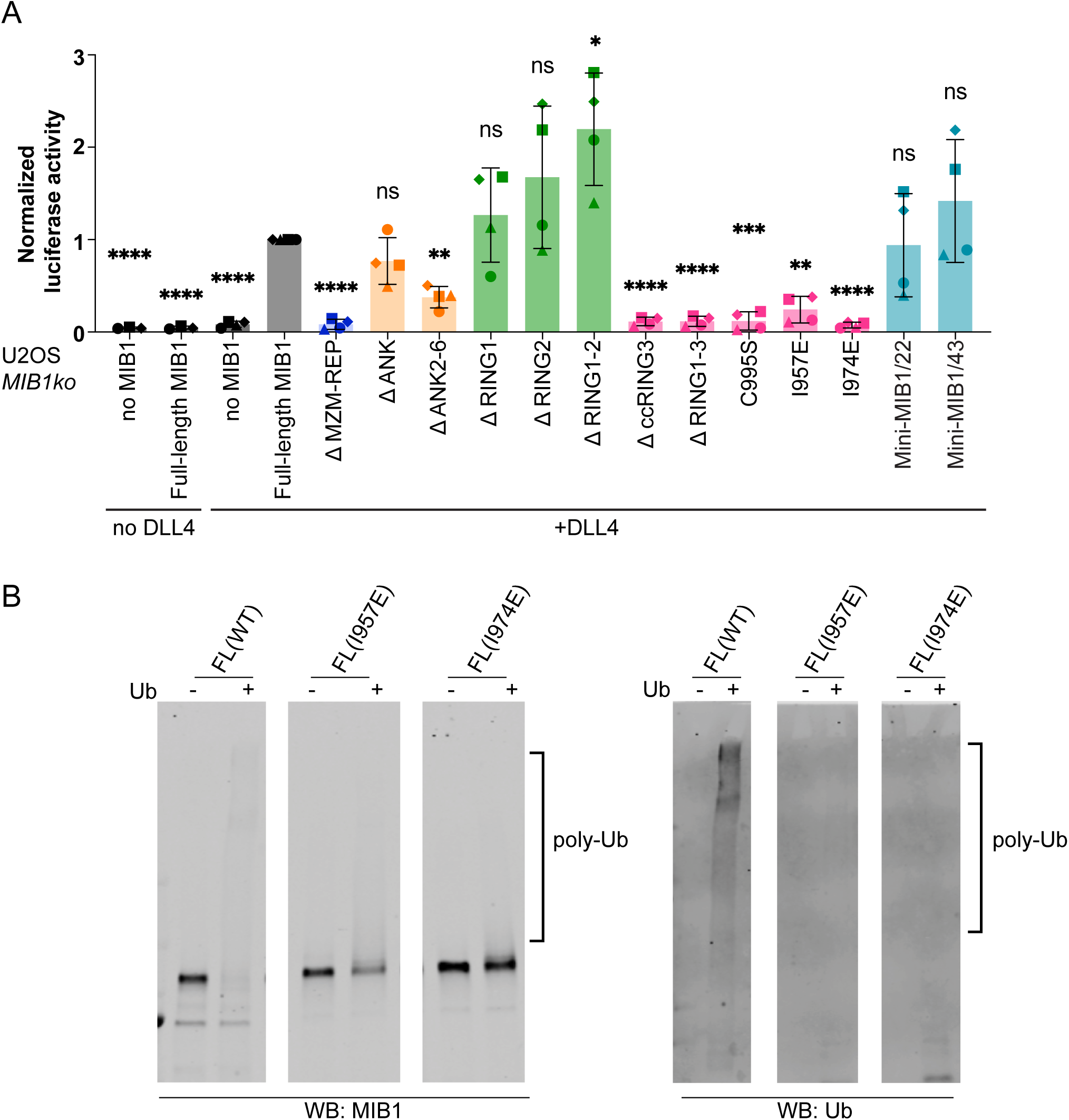
Reporter gene and ubiquitination assays testing the activity of MIB1 variants. A, Reporter gene assay. Activation of U2OS-Notch1-Gal4 receiver cells by U2OS MIB1 knockout cells stably transfected with Full-length MIB1 alone, DLL4 alone or DLL4 and wild-type (Full-length) or mutant/truncated forms of MIB1. All results were normalized to the activity with U2OS MIB1 knockout cells stably transfected with DLL4 and wild-type MIB1. Error bars represent standard deviation over n ≥ 3 independent repeats, and p-values were determined using a one sample Wilcoxon signed rank test. ns: p>0.05, *, p<0.05, **, p<0.01; ***, p<0.001, ****, p<0.0001. B, Ubiquitination assay. Wild-type (FL(WT)), I957E, and I974E forms of MIB1 were incubated with ubiquitin, E1, and E2. Protein products were separated by SDS-PAGE on 4-20% gradient gels. Products were detected by Western blot with an anti-MIB1 antibody (left), or an anti-Ubiquitin antibody (right).

Remarkably, despite the conservation of the concave face of the ANK domain (Supplementary Fig. 3B), deletion of the entire ANK domain results only in a small reduction of Notch reporter activity, an effect mimicked by a smaller internal deletion within the ANK domain. In addition, deletion of either RING1 or RING2 has no significant effect on signaling, whereas deletion of both RING1 and RING2 increases the reporter signal, suggesting that these two repeats together might perform a minor autoinhibitory role.

### Effect of dimer disrupting mutations on MIB1 activity

The isolated ccRING3 region forms a homodimer, as judged by multiple angle light scattering (SEC-MALS; Supplementary Fig. 5A). In contrast, SEC-MALS revealed that the isolated, purified ANK domain is a monomer (Supplementary Fig. 5B), that ANK-RING1-2 is predominantly monomeric (Supplementary Fig. 5C), and that ANK-RING1-3 forms a homodimer (Supplementary Fig. 5D). The MZM/REP region is monomeric in isolation, as judged by small angle X-ray scattering^17^, and Alphafold multimer^34^ also predicts that the isolated ccRING3 region dimerizes using the interface seen in the crystal structure of the UbcH5B-ccRING3 complex (Fig. 2). Together, these data indicate that dimerization of MIB1 relies primarily on the ccRING3 region.

To test the functional importance of dimerization of the cc-RING3 region of MIB1, we introduced an I957E mutation to disrupt hydrophobic packing of the coiled coil, or an I974E mutation within the core RING3 domain (Supplementary Fig. 6A). We purified each mutated ccRING3 protein to homogeneity (Supplementary Fig. 6B) and confirmed that each mutation converted ccRING3 from a dimer to a monomer, as judged using SEC-MALS (Supplementary Fig. 6C). When introduced into full-length MIB1, the I974E mutation did not result in a detectable increase in signaling activity compared to the negative control in the reporter assay, whereas I957E resulted in partial, but incomplete rescue of signaling activity (Fig. 4A). A C995S mutation in the RING3 domain, which disables the RING domain of other E3 ligases and inactivates MIB1 in other contexts,^12,25^ also eliminates MIB1 activity in the reporter assay (Fig. 4A).

We conducted an autoubiquitination assay with purified full-length wild-type (FL(WT)) MIB1 protein and compared it with the activity of the purified I957E and I974E mutants using comparable amounts of MIB1 protein (Fig. 4B). The autoubiquitination activity of the FL(WT), I957E, and I974E proteins closely tracked their relative signaling activity in the reporter assay, with WT MIB1 exhibiting extensive autoubiquitination, the I957E mutant partial autoubiquitination, and the I974E mutant negligible autoubiquitination (Fig. 4B). Together, the reporter and ubiquitination data show that the MIB1 dimer is more active than the monomer, but that dimerization is not an absolute prerequisite for activity.

### Construction of a functional mini-MIB1 without the ANK-RING1-2 region

Because the ANK, RING1, and RING2 domains are not essential for MIB1 activity in the reporter gene assay, we tested whether tethering MZM/REP to the ccRING3 region of MIB1 with a flexible linker is sufficient to support DLL4-induced Notch signaling in the co-culture assay. We connected the MZM/REP and ccRING3 elements with two different-length linkers, one of 22 residues (Mini-MIB1/22) and the other of 43 residues (Mini-MIB1/43). Both Mini-MIB1 proteins induced a signaling response indistinguishable from wild-type (Full-length) MIB1 in the reporter gene assay (Fig. 4A), confirming that the ANK, RING1, and RING2 domains are dispensable for the function of MIB1 in Notch signal-sending cells.

## Discussion

This work reports a structure-function analysis of the ANK and RING regions of human MIB1 in Notch-Delta signaling. Previous studies have shown that the N-terminal MZM and REP domains bind the cytoplasmic tails of Notch ligands and defined the structural basis for ligand tail recognition.^17^ Early studies of the zebrafish and fly Mib proteins also implicated the third RING domain as most critical for function in Notch signaling.^12,36,37^

The contributions of the ANK domain and the other two RING domains in the function of MIB as a potentiator of Notch ligands, however, have been less clear. Work in flies showed that deletion of all three RING domains had dominant negative activity in Notch signaling like that seen when only RING3 was deleted, and forced expression of a variant containing only the ANK and RING domains did not show a phenotype in the same assay.^36^ Others also showed that proteins lacking only MZM/REP or the ANK domains had strong autoubiquitination activity, whereas deletion of RING3 or all three RING domains suppressed ubiquitination of Delta in cellular assays.^12,36,37^

Our studies with purified proteins showed that the ccRING3 region of MIB1 was absolutely required for E3 ligase activity when UbcH5B is used as an E2 subunit. In contrast, there was no detectable ubiquitination activity in proteins that include RING1 and RING2 but lack ccRING3. Moreover, a MIB1 deletant lacking RING1 and RING2 was completely able to substitute for wild type MIB1 in enabling DLL4 to stimulate Notch reporter activity in a cellular co-culture assay, establishing that RING1 and RING2 are not required for MIB1 function in Notch signal induction.

Most strikingly, deletion of roughly half of the protein, including removal of the ANK, RING1, and RING2 domains, had no appreciable effect on the ability of MIB1 to support the activity of DLL4 as a functional Notch ligand in a co-culture assay. This finding established that the minimal elements required to potentiate DLL4 as a Notch ligand were the MZM-REP region and the ccRING3 domain, connected by a flexible tether, which approaches the activity of the full-length protein in Notch signaling assays (Fig. 5). The dispensability of the ANK, RING1 and RING2 domains was unexpected because these domains are conserved in all metazoan MIB proteins. Moreover, the high conservation of the concave surface of the ANK domain suggests that it serves as a binding interface for a partner protein to execute one of its biological functions. Testing whether the ANK domain (as well as the RNIG1 and RING2 domains) required for other cellular roles of MIB1, such as internalization of the RYK tyrosine kinase and centriolar satellite formation, should be fertile ground for future studies.

**Fig. 5.**
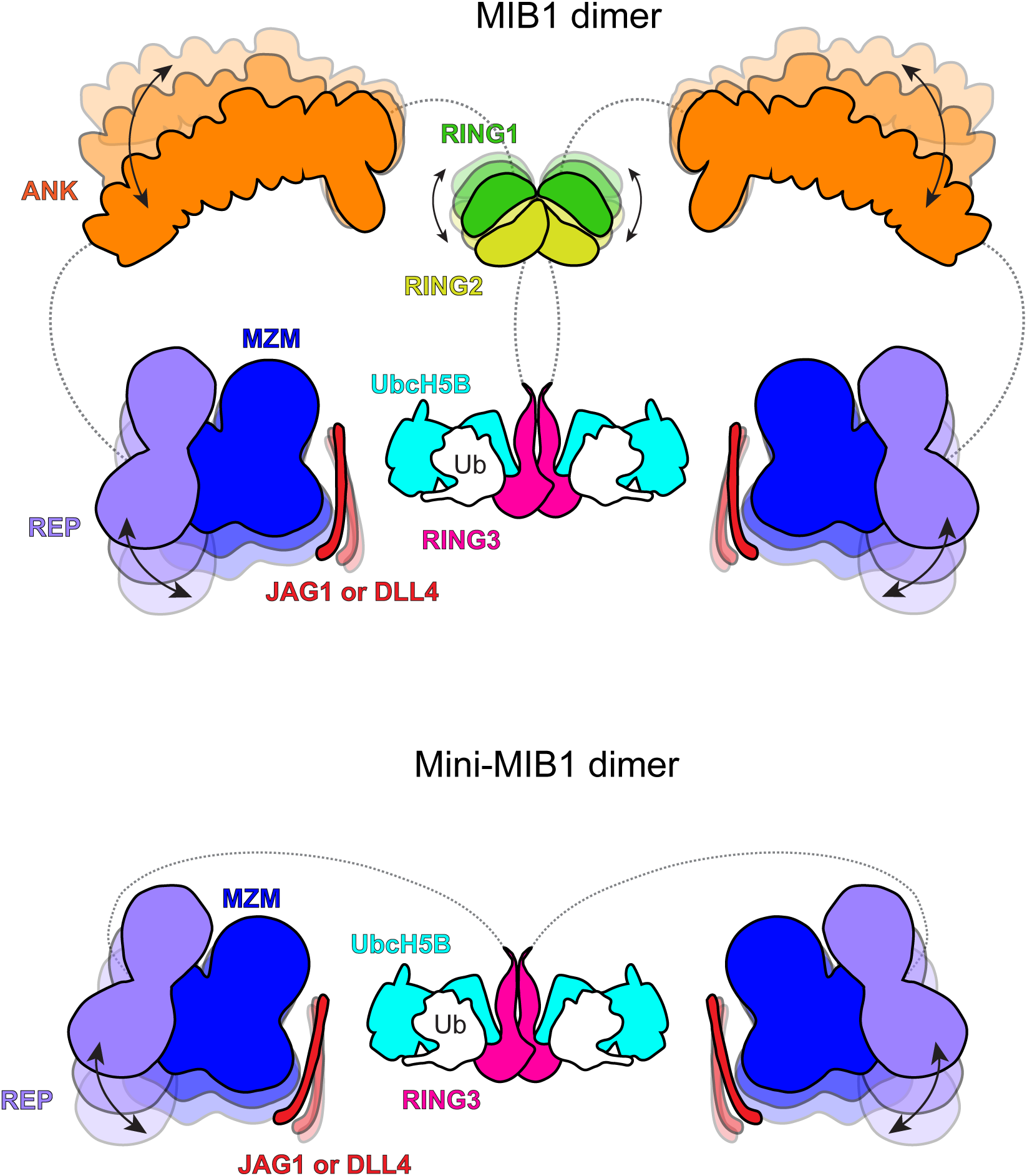
Schematic models of MIB1 (top) and Mini-MIB1 (bottom) ligase complexes. The N-terminal MZM-REP region of MIB1 recognizes a NOTCH ligand (*e.g.* JAG1 or DLL4), and the ccRING3 domain engages UbcH5B-Ub to promote ubiquitin (Ub) transfer. The linkers connecting MZM-REP to ANK and ANK to the RING1-2 region are flexible (schematically illustrated by lighter shading of domains and arrows to indicate movement) and not required for the signal activation function of NOTCH ligands.

## Author contributions

S.C.B. and R.C. conceived the project with input from D.S. and O.G. S.C.B. acquired funding. R.C., S.C.B., O.G., D.S., L.T., J.C.A., H.Z., H.J., P.C. designed experiments. R.C. and O.G. performed experiments. R.C., L.T. and O.G. analyzed data. All authors assisted with data analysis and interpretation. S.C.B. and R.C. wrote the manuscript with input from all authors. All authors provided feedback and agreed on the final manuscript.

## Acknowledgments

We thank all members of the Blacklow laboratory for helpful discussions and encouragement. This work was supported by NIH award 1R35 CA220340 (to S.C.B.), R01 CA74305 (to P.A.C.), and the Ludwig Center at Harvard (to J.C.A.). This research was also supported by grant No 2017245 from the United States-Israel Binational Science Foundation (BSF) (to S.C.B. and D.S.).

## Methods

### Plasmid construction

ANK-RING1-3 (residues 417-1006) of hMIB1 was cloned into the pGood6p vector with an N-terminal GST tag followed by a 3C cleavage site. RING1-3 (residues 813-1006), RING1-2 (residues 813-919), ccRING3 (residues 936-1006) were cloned into the ptd68 vector with an N-terminal HIS-SUMO tag. ANK-RING1-2 (residues 409-919), ANK (residues 409-794) and UbcH5B-GSlinker-ccRING3 were cloned into a pET-21a vector with an N-terminal His tag followed by a 3C cleavage site. Full length human MIB1 was cloned into a pcDNA3.1/hygro(+) vector with an N-terminal Flag tag. The mutations of full length MIB1 were made using site-directed mutagenesis. All MIB1 domain truncations were generated using overlap PCR. mMib1-Flag was purchased from Addgene (catalog #37116).

### Protein expression and purification

Recombinant ANK, ANK-RING1-2, ANK-RING1-3, RING1-3, RING1-2, ccRING3, and UbcH5B-GSlinker-ccRING3 proteins were produced in *E. coli* BL21(DE3) cells. Expression was induced with 0.2 mM isopropyl-1-thio-D-galactopyranoside (IPTG), and cells were grown overnight at 16°C. 100 μM ZnCl_2_ was added at the time of induction when at least one RING domain was present in the expressed construct. Cells were harvested by centrifugation and resuspended in lysis buffer. For cells expressing His-tagged proteins, the lysis buffer was 20 mM Tris HCl, pH 7.6, containing 150 mM NaCl, 20 mM Imidazole, and 2 mM TCEP. For cells expressing proteins that were not His-tagged, the lysis buffer was 20 mM Tris HCl, pH 7.6, containing 150 mM NaCl, and 2 mM TCEP (without Imidazole). After cells were lysed by sonication, the lysate was centrifuged to remove cell debris, and the supernatant was collected. Recombinant His-tagged proteins were affinity purified using Ni-NTA beads and GST-tagged proteins were affinity purified using glutathione beads. Each affinity captured protein was washed with 20 column volumes of lysis buffer before proteolytically releasing the tag by on-column cleavage. His-SUMO tagged proteins were released with ULP1 protease; GST-tagged and His-tagged proteins were released using 3C protease. The released MIB1 proteins were recovered, and further purified on a size exclusion column (Superdex 200 10/300 GL or Superdex 75 10/300 GL, depending on the construct) in 20 mM Tris HCl buffer, pH 7.6, containing 100 mM NaCl, and 2 mM TCEP.

Full length human MIB1 WT, I957E, I974E proteins and murine MIB1 protein were expressed in Expi293F cells. Cells were grown in Expi293 media to a density of 3 x 10^6^ cells/ml and then transfected with 1.0 mg DNA/L of culture using the FectroPro transfection reagent (Polyplus) at a 1:1 DNA/FectroPro ratio. 24 hours after transfection, 45% D-(+)-Glucose solution (Sigma-Aldrich, 10 mL per L of culture) and 3 mM valproic acid sodium salt (Sigma-Aldrich) were added to the cells to enhance protein expression. The cells were cultured for an additional 24 hours before harvesting by centrifugation. Cells were then resuspended in lysis buffer. For hMIB1 and mutants, the lysis buffer was 20 mM HEPES pH 7.5, containing 500 mM NaCl, 10% Glycerol, 0.02% Tween 20, and 2 mM TCEP. For mMIB1, the lysis buffer was 20 mM HEPES pH 7.5, containing 500 mM NaCl, 10% Glucose, 100 mM L-Arginine, 10 μM ZnCl_2_, and 0.5 mM TCEP. After sonication, the lysate was centrifuged to remove cell debris, and the supernatant was collected. Recombinant Flag-tagged proteins were affinity purified using anti-Flag resin, washed with 20 column volumes of lysis buffer, and eluted with lysis buffer containing Flag peptide (0.2 mg/mL). Eluted WT and mutant MIB1 proteins were concentrated and used in the ubiquitination activity assay. mMIB1 was concentrated and further purified using Superdex 6 10/300 GL in 20 mM HEPES pH 7.5, containing 500 mM NaCl, 10% Glucose, 100 mM L-Arginine, 10 μM ZnCl_2_, and 0.5 mM TCEP. Protein purity was assessed by SDS-PAGE followed by Coomassie blue staining. Protein concentrations were determined by UV absorbance at 280 nm.

### Crystallization, data collection and structure determination

UbcH5B-GSlinker-ccRING3 was concentrated to 20 mg/ml in 20 mM Tris HCl, pH 7.6 buffer containing 100 mM NaCl, and 2 mM TCEP. Crystals were grown in sitting drops at 18°C by mixing equal volumes of protein and a reservoir solution containing 4% (v/v) Tacsimate pH 6.0, and 12% (w/v) polyethylene glycol 3,350. Microseeding was then performed to optimize growth of single crystals. Crystals were cryoprotected in reservoir solution supplemented with 25% (v/v) glycerol, and flash frozen in liquid nitrogen for shipment and data collection at Advanced Photon Source NE-CAT beamlines 24 ID-C and ID-E. UbcH5B (PDB ID code: 2ESK) and an AlphaFold predicted^38^ ccRING3 model were used as search models for molecular replacement in Phaser.^39^ Model building and refinement were carried out with the programs COOT^40^ and PHENIX.^41^ Data collection and structural refinement statistics are reported in Table S1.

ANK was concentrated to 15 mg/mL in 20 mM Tris HCl, pH 7.6 buffer containing 100 mM NaCl, and 2 mM TCEP. Crystals were grown in sitting drops at 18° C by mixing equal volumes of protein and a reservoir solution containing 0.1 M Sodium acetate trihydrate, and 1.0 M Ammonium tartrate (dibasic) at pH 4.6. Crystals were cryoprotected in reservoir solution supplemented with 25% (v/v) glycerol, and flash frozen in liquid nitrogen for shipment and data collection at Advanced Photon Source NE-CAT beamlines 24 ID-C and ID-E. Diffraction images were indexed, integrated, and merged using HKL2000.^42^ The phase was determined using the ankyrin domain of a DARPIN-erythropoetin receptor complex (PDB ID code:6MOL) as a search model for molecular replacement in Phaser.^39^ Model building and refinement were carried out with the programs COOT^40^ and PHENIX.^41^ Data collection and structural refinement statistics are reported in Table S2.

### Negative-stain electron microscopy

Carbon-coated copper grids (Electron Microscopy Sciences, #CF400-Cu) were glow discharged at 30 mA for 30 s. A 4 μL aliquot of a mMIB1 full length sample (0.02 mg/mL) was applied to the grid. After incubating for 1 min, the grid was washed twice with 1.5% Uranyl formate followed by staining with 1.5% Uranyl formate for 2 min. The grids were imaged using a 120 kV Tecnai T12 (Thermo Fisher Scientific) microscope. Images were recorded using an Ultrascan 895 CCD camera (Gatan).

### In vitro Ubiquitination assays

Ubiquitination was performed by mixing E1 enzyme (50 nM), UbcH5B (500 nM), a MIB1 variant (1 μM), and Ubiquitin (5 μM) in a reaction buffer containing 25 mM HEPES pH 7.5, 100 mM NaCl, 5 mM MgCl_2_, 5 mM ATP, and 2 mM TCEP at 37°C for 3 h. SDS-loading buffer was added directly to each tube to terminate the reaction. 10 μL of each sample was subjected to SDS-PAGE and analyzed by staining with Coomassie blue dye. Duplicate samples (0.5 μL) were subjected to SDS-PAGE and analyzed by western blot with an anti-Ubiquitin antibody (Santa Cruz Biotechnology, sc-8017). When using full length MIB1 protein, a western blot with anti-MIB1 (Abcam, ab124929) antibody was also used to analyze the ubiquitin reaction.

### Generation of cells: CRISPR/Cas9 Knockout of MIB1 und lentiviral transduction of DLL4 and MIB1 in U2OS cells

sgRNA sequences targeting bp 171-193 in the first exon of human *MIB1* were designed using the chopchop online tool (https://chopchop.cbu.uib.no) and subcloned into the plasmid vector PX458. The forward sgRNA sequence was 5’-CACCGTGCCAACTACCGCTGCTCCG-3’ and the sequence for the reverse sgRNA was 5’-AAACCGGAGCAGCGGTAGTTGGCAC-3’. U2OS cells were transfected in 6-well plates with 4 μg sgRNA-PX458 and 10 μl lipofectamine. 48 hours after transfection, single GFP-positive cells were sorted by FACS. Each clone was grown until it was confluent on a 10 cm dish. MIB1 knockout was confirmed by TOPO cloning of the amplified genomic sequence and by the loss of MIB1 signal on western blot with antibodies for the N- and C-terminal portions of MIB1 (Abcam, ab124929; Millipore Sigma, M5948).

Wild-type DLL4 and wild-type or mutated forms of MIB1 were introduced into MIB1 knockout U2OS cells using lentiviral transduction in order to test the activities of different MIB1 variants in reporter gene activity assays. MIB1 and DLL4 proteins were inserted into the vector pLVX. All MIB1 variants were fused to mNeonGreen at their N-termini and DLL4 was tagged with mTurquoise2 at its C-terminus.

### Co-culture luciferase reporter assays

Receiver (Notch-expressing) cells were U2OS-Notch1-Gal4 cells^43^ and sender cells were the MIB1 knockout cells lentivirally transfected as described above. All cells were cultured in Dulbecco’s Modified Eagle’s Medium (DMEM) and grown in an environment of 5% CO2 at 37°C. Receiver cells were transfected (Lipofectamine 3000) with a UAS-firefly luciferase reporter (350 ng) and pRL-SV40 Renilla luciferase (10 ng), in addition to a H2B-Cerulean plasmid (10 ng) to test transfection efficiency. Receiver cells were transferred onto plates containing the sender cells 24 h after transfection. Cells were lysed 24 h after the start of co-culture. Firefly and Renilla luciferase activities were measured using a luminometer (Promega GloMax(R) Navigator with Dual Injectors). Relative Notch activity was calculated from the ratio of luciferase to Renilla signal, normalizing to the value for MIB1 knockout sender cells lentivirally transduced with wild-type DLL4 and MIB1.

### Flow cytometry

The amount of expressed MIB1-GFP was analyzed by flow cytometry for each variant. Cells were harvested from tissue culture plates with trypsin/EDTA in phosphate buffered saline containing fetal bovine serum (1% v/v). All analyses were performed on the same day with the same parameters, using a 488 nm laser on a CytoFLEX S Flow Cytometer (Beckman Instruments).

### Statistical analysis

Statistical analysis was performed using GraphPad Prism version 10 (GraphPad). Statistical details are indicated in the figure legend along with the value of n. Sample distribution and normality tests were performed for the data set and significance was determined using a one sample Wilcoxon signed rank test.

## Supplementary Figure Legends

**Supplementary Fig. 1. In vitro ubiquitination assay.** Western blot analysis of *in vitro* auto-ubiquitylation assays with ANK-RING1-3 (A) and RING1-3 (B). ANK-RING1-3 and RING1-3 only show activity in the presence of Ub, E1 and E2.

**Supplementary Fig. 2. Sequence alignment of RING3 from MIB1 with RING domains from other E3 ligases**. Conserved positions are highlighted (red). Asterisks indicate residues interacting with E2 proteins in x-ray crystal structures of E2-E3 complexes.

**Supplementary Fig. 3. Crystal packing in the x-ray structure and conservation analysis of the ANK domain of MIB1.** Two copies of the ankyrin domain are shown in cartoon representation. The interface between the extended helices of ankyrin repeat nine is indicated with a dotted oval. B. Consurf analysis^44^ of the ankyrin repeat domain of MIB1, colored on a sliding scale from teal (poorly conserved) to maroon (highly conserved). Note the high conservation of the concave face and the poor conservation of the convex face.

**Supplementary Fig. 4. MIB1 variants tested and flow cytometry analysis of expression**. A, Schematic of mNeonGreen-MIB1 variants tested in the reporter gene assay. Internally deleted regions are indicated with a dotted line, and point mutations are indicated with stars. B, Flow cytometry analysis of mNeonGreen-MIB1 protein abundance in cell lines stably expressing different MIB1 variants. Histograms of the channel detecting mNeonGreen are shown.

**Supplementary Fig. 5. SEC-MALS analysis of ccRING3 (A), ANK (B), ANK-RING1-2 (C) and ANK-RING1-3 (D).** The traces show that ccRING3 and ANK-RING1-3 are dimeric, whereas ANK is monomeric and ANK-RING1-2 is predominantly monomeric.

**Supplementary Fig. 6. Identification, purification, and analysis of dimer-disrupting mutants of ccRING3.** A, Cartoon representation showing dimerization contacts in the coiled coil (left) and RING3 (right) domains. Interfacial hydrophobic residues are labelled, and mutated isoleucine residues are boxed in red. B, SDS-PAGE analysis of purified ccRING3 I957E and I974E mutants. Proteins were visualized by staining with Coomassie blue. C, SEC-MALS comparison of wild-type (black trace), I957E (red trace), and I974E (green trace) forms of cc-RING3.

**Table S1.** Data collection and refinement statistics.

|  | UbcH5B-ccRING3 |
| --- | --- |
| **Data collection and processing** |  |
| Wavelength (Å) | 0.9792 |
| Resolution range (Å) | 37.16 - 2.40 (2.49 - 2.40) |
| Space group (Å, °) | P 1 21 1 |
| Unit cell | 28.999, 65.12, 135.93, 90, 93.03, 90 |
| Total reflections | 1714570 |
| Unique reflections | 18194 (1655) |
| Multiplicity | 3.0 (3.0) |
| Completeness (%) | 91.35 (84.78) |
| Mean I/sigma(I) | 16.86 (1.8) |
| Wilson B-factor | 45.76 |
| R-merge | 0.086 (0.443) |
| R-meas | 0.104 (0.531) |
| R-pim | 0.057 (0.288) |
| CC1/2 | 0.990 (0.903) |
| **Refinement** |  |
| Reflections used in refinement | 18183 (1655) |
| Reflections used for R-free | 938 (84) |
| R-work | 0.2134 (0.3100) |
| R-free | 0.2443 (0.3021) |
| Model composition |  |
| Number of non-hydrogen atoms | 3435 |
| Macromolecules | 3391 |
| Ligands | 5 |
| Solvent | 39 |
| Protein residues | 424 |
| *B* factors (Å<sup>2</sup>) |  |
| Macromolecules | 64.57 |
| Ligand | 60.01 |
| Solvent | 55.01 |
| R.m.s. deviations |  |
| Bond lengths (Å) | 0.007 |
| Bond angles (°) | 0.96 |
| Ramachandran plot |  |
| Favored (%) | 97.60 |
| Allowed (%) | 2.40 |
| Outliers (%) | 0.00 |
| Rotamer outliers (%) | 0.52 |
| Clashscore | 6.34 |
| Number of TLS groups | 13 |

Statistics for the highest-resolution shell are shown in parentheses.

**Table S2.**
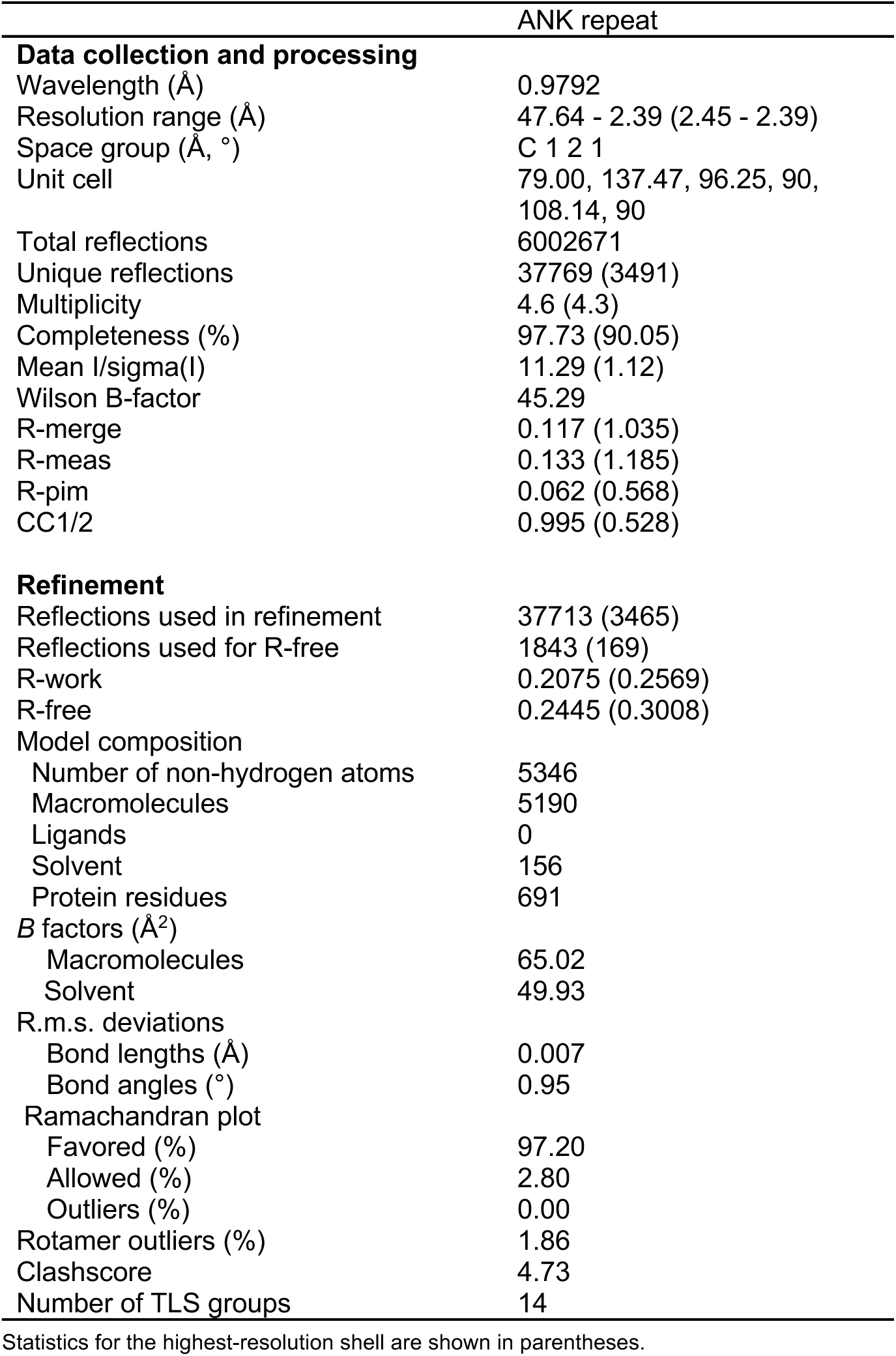
Data collection and refinement statistics.

## References

1. Artavanis-Tsakonas, S., Rand, M.D., and Lake, R.J. (1999). Notch signaling: cell fate control and signal integration in development. Science 284, 770–776. 10.1126/science.284.5415.770.

2. Lewis, J. (1998). Notch signalling and the control of cell fate choices in vertebrates. Semin Cell Dev Biol 9, 583–589. 10.1006/scdb.1998.0266.

3. Siebel, C., and Lendahl, U. (2017). Notch Signaling in Development, Tissue Homeostasis, and Disease. Physiol Rev 97, 1235–1294. 10.1152/physrev.00005.2017.

4. Schroeter, E.H., Kisslinger, J.A., and Kopan, R. (1998). Notch-1 signalling requires ligand-induced proteolytic release of intracellular domain. Nature 393, 382–386. 10.1038/30756.

5. G Struhl, I.G. (2001). Presenilin-mediated transmembrane cleavage is required for Notch signal transduction in Drosophila. Proc Natl Acad Sci U S A 98, 229–234.

6. F Oswald, B.T., T Dobner, S Bourteele, U Kostezka, G Adler, S Liptay, R M Schmid (2001). p300 acts as a transcriptional coactivator for mammalian Notch-1. Mol Cell Biol 21, 7761–7774.

7. Nam, Y., Sliz, P., Song, L., Aster, J.C., and Blacklow, S.C. (2006). Structural basis for cooperativity in recruitment of MAML coactivators to Notch transcription complexes. Cell 124, 973–983. 10.1016/j.cell.2005.12.037.

8. Wilson, J.J., and Kovall, R.A. (2006). Crystal structure of the CSL-Notch-Mastermind ternary complex bound to DNA. Cell 124, 985–996. 10.1016/j.cell.2006.01.035.

9. Kelly L Arned, M.H., Debbie G McArthur, Ma Xenia G Ilagan, Jon C Aster, Raphael Kopan, Stephen C Blacklow (2010). Structural and mechanistic insights into cooperative assembly of dimeric Notch transcription complexes. Nat Struct Mol Biol. 17, 1312–1317.

10. Musse, A.A., Meloty-Kapella, L., and Weinmaster, G. (2012). Notch ligand endocytosis: mechanistic basis of signaling activity. Semin Cell Dev Biol 23, 429–436. 10.1016/j.semcdb.2012.01.011.

11. Haddon, C., Jiang, Y.J., Smithers, L., and Lewis, J. (1998). Delta-Notch signalling and the paderning of sensory cell differentiation in the zebrafish ear: evidence from the mind bomb mutant. Development 125, 4637–4644. 10.1242/dev.125.23.4637.

12. Itoh, M., Kim, C.H., Palardy, G., Oda, T., Jiang, Y.J., Maust, D., Yeo, S.Y., Lorick, K., Wright, G.J., Ariza-McNaughton, L., et al. (2003). Mind bomb is a ubiquitin ligase that is essential for efficient activation of Notch signaling by Delta. Dev Cell 4, 67–82. 10.1016/s1534-5807(02)00409-4.

13. EricC.Lai, G.l.D., ChrisKintner,andGerald M.Rubin (2001). Drosophila Neuralized Is a Ubiquitin Ligase that Promotes the Internalization and Degradation of Delta. Dev Cell 1, 783–794.

14. Daskalaki, A., Shalaby, N.A., Kux, K., Tsoumpekos, G., Tsibidis, G.D., Muskavitch, M.A., and Delidakis, C. (2011). Distinct intracellular motifs of Delta mediate its ubiquitylation and activation by Mindbomb1 and Neuralized. J Cell Biol 195, 1017–1031. 10.1083/jcb.201105166.

15. Roland Le Borgne, S.R., Sophie Hamel, François Schweisguth (2005). Two distinct E3 ubiquitin ligases have complementary functions in the regulation of delta and serrate signaling in Drosophila. PloS Biol. 3, e96.

16. Bon-Kyoung Koo, K.-J.Y., Kyeong-Won Yoo, Hyoung-Soo Lim, Ran Song, Ju-Hoon So,, and Cheol-Hee Kim, a.Y.-Y.K. (2005). Mind Bomb-2 Is an E3 Ligase for Notch Ligand. THE JOURNAL OF BIOLOGICAL CHEMISTRY 280, 22335–22342.

17. McMillan, B.J., Schnute, B., Ohlenhard, N., Zimmerman, B., Miles, L., Beglova, N., Klein, T., and Blacklow, S.C. (2015). A tail of two sites: a bipartite mechanism for recognition of notch ligands by mind bomb E3 ligases. Mol Cell 57, 912–924. 10.1016/j.molcel.2015.01.019.

18. Koo, B.K., Yoon, M.J., Yoon, K.J., Im, S.K., Kim, Y.Y., Kim, C.H., Suh, P.G., Jan, Y.N., and Kong, Y.Y. (2007). An obligatory role of mind bomb-1 in notch signaling of mammalian development. PLoS One 2, e1221. 10.1371/journal.pone.0001221.

19. Barsi, J.C., Rajendra, R., Wu, J.I., and Artzt, K. (2005). Mind bomb1 is a ubiquitin ligase essential for mouse embryonic development and Notch signaling. Mech Dev 122, 1106–1117. 10.1016/j.mod.2005.06.005.

20. Koo, B.K., Lim, H.S., Song, R., Yoon, M.J., Yoon, K.J., Moon, J.S., Kim, Y.W., Kwon, M.C., Yoo, K.W., Kong, M.P., et al. (2005). Mind bomb 1 is essential for generating functional Notch ligands to activate Notch. Development 132, 3459–3470. 10.1242/dev.01922.

21. Hyun-Woo, J., and Ji-Hoon K., J.-Y., Sang-JunHa2,Young-YunKong1* (2012). Mind Bomb-1 in Dendritic Cells Is Specifically Required for Notch-mediated T Helper Type2 Differentiation. PLoS ONE 7, e36359.

22. Ran, S. (2008). Mind bomb 1 in the lymphopoietic niches is essential for T and marginal zone B cell development. J Exp Med. 205, 2525–2536.

23. Yoon, M.J. (2008). Mind bomb-1 is essential for intraembryonic hematopoiesis in the aortic endothelium and the subaortic patches. Mol Cell Biol. 28, 4794–4804.

24. Luxan, G., Casanova, J.C., Martinez-Poveda, B., Prados, B., D’Amato, G., MacGrogan, D., Gonzalez-Rajal, A., Dobarro, D., Torroja, C., Martinez, F., et al. (2013). Mutations in the NOTCH pathway regulator MIB1 cause lem ventricular noncompaction cardiomyopathy. Nat Med 19, 193–201. 10.1038/nm.3046.

25. Berndt, J.D., Aoyagi, A., Yang, P., Anastas, J.N., Tang, L., and Moon, R.T. (2011). Mindbomb 1, an E3 ubiquitin ligase, forms a complex with RYK to activate Wnt/beta-catenin signaling. J Cell Biol 194, 737–750. 10.1083/jcb.201107021.

26. Wang, L., Lee, K., Malonis, R., Sanchez, I., and Dynlacht, B.D. (2016). Tethering of an E3 ligase by PCM1 regulates the abundance of centrosomal KIAA0586/Talpid3 and promotes ciliogenesis. Elife 5. 10.7554/eLife.12950.

27. Villusen, B.H. (2013). A new cellular stress response that triggers centriolar satellite reorganization and ciliogenesis. EMBO J. 32, 3029–3040.

28. Bauer, M. (2019). The E3 Ubiquitin Ligase Mind Bomb 1 Controls Adenovirus Genome Release at the Nuclear Pore Complex. Cell Rep. 29, 3785–3795.

29. Liu, L.J., Liu, T.T., Ran, Y., Li, Y., Zhang, X.D., Shu, H.B., and Wang, Y.Y. (2012). The E3 ubiquitin ligase MIB1 negatively regulates basal IkappaBalpha level and modulates NF-kappaB activation. Cell Res 22, 603–606. 10.1038/cr.2011.199.

30. Zhang, L., and Gallagher, P.J. (2009). Mind bomb 1 regulation of cFLIP interactions. Am J Physiol Cell Physiol 297, C1275–1283. 10.1152/ajpcell.00214.2009.

31. Dou, H., Buetow, L., Sibbet, G.J., Cameron, K., and Huang, D.T. (2012). BIRC7-E2 ubiquitin conjugate structure reveals the mechanism of ubiquitin transfer by a RING dimer. Nat Struct Mol Biol 19, 876–883. 10.1038/nsmb.2379.

32. Mace, P.D., Linke, K., Feltham, R., Schumacher, F.R., Smith, C.A., Vaux, D.L., Silke, J., and Day, C.L. (2008). Structures of the cIAP2 RING domain reveal conformational changes associated with ubiquitin-conjugating enzyme (E2) recruitment. J Biol Chem 283, 31633–31640. 10.1074/jbc.M804753200.

33. Magnussen, H.M., Ahmed, S.F., Sibbet, G.J., Hristova, V.A., Nomura, K., Hock, A.K., Archibald, L.J., Jamieson, A.G., Fushman, D., Vousden, K.H., et al. (2020). Structural basis for DNA damage-induced phosphoregulation of MDM2 RING domain. Nat Commun 11, 2094. 10.1038/s41467-020-15783-y.

34. Mirdita, M., Schütze, K., Moriwaki, Y., Heo, L., Ovchinnikov, S., and Steinegger, M. (2022). ColabFold: making protein folding accessible to all. Nature Methods 19, 679–682.

35. Aster, J.C., Xu, L., Karnell, F.G., Patriub, V., Pui, J.C., and Pear, W.S. (2000). Essential roles for ankyrin repeat and transactivation domains in induction of T-cell leukemia by notch1. Mol Cell Biol 20, 7505–7515. 10.1128/MCB.20.20.7505-7515.2000.

36. Lai, E.C., Roegiers, F., Qin, X., Jan, Y.N., and Rubin, G.M. (2005). The ubiquitin ligase Drosophila Mind bomb promotes Notch signaling by regulating the localization and activity of Serrate and Delta. Development 132, 2319–2332. 10.1242/dev.01825.

37. Chen, W., and Casey Corliss, D. (2004). Three modules of zebrafish Mind bomb work cooperatively to promote Delta ubiquitination and endocytosis. Dev Biol 267, 361–373. 10.1016/j.ydbio.2003.11.010.

38. Jumper, J., Evans, R., Pritzel, A., Green, T., Figurnov, M., Ronneberger, O., Tunyasuvunakool, K., Bates, R., Zidek, A., Potapenko, A., et al. (2021). Highly accurate protein structure prediction with AlphaFold. Nature 596, 583–589. 10.1038/s41586-021-03819-2.

39. McCoy, A.J. (2007). Solving structures of protein complexes by molecular replacement with Phaser. Acta Crystallogr D Biol Crystallogr 63, 32–41. 10.1107/S0907444906045975.

40. Emsley, P., and Cowtan, K. (2004). Coot: model-building tools for molecular graphics. Acta Crystallogr D Biol Crystallogr 60, 2126–2132. 10.1107/S0907444904019158.

41. Adams, P.D., Afonine, P.V., Bunkoczi, G., Chen, V.B., Davis, I.W., Echols, N., Headd, J.J., Hung, L.W., Kapral, G.J., Grosse-Kunstleve, R.W., et al. (2010). PHENIX: a comprehensive Python-based system for macromolecular structure solution. Acta Crystallogr D Biol Crystallogr 66, 213–221. 10.1107/S0907444909052925.

42. Otwinowski, Z., and Minor, W. (1997). Processing of X-ray diffraction data collected in oscillation mode. Methods Enzymol 276, 307–326. 10.1016/S0076-6879(97)76066-X.

43. Gordon, W.R., Vardar-Ulu, D., L’Heureux, S., Ashworth, T., Malecki, M.J., Sanchez-Irizarry, C., McArthur, D.G., Histen, G., Mitchell, J.L., Aster, J.C., and Blacklow, S.C. (2009). Effects of S1 cleavage on the structure, surface export, and signaling activity of human Notch1 and Notch2. PLoS One 4, e6613. 10.1371/journal.pone.0006613.

44. Ashkenazy, H., Abadi, S., Martz, E., Chay, O., Mayrose, I., Pupko, T., and Ben-Tal, N. (2016). ConSurf 2016: an improved methodology to estimate and visualize evolutionary conservation in macromolecules. Nucleic Acids Res 44, W344–350. 10.1093/nar/gkw408.

