## Supplementary_figures for "Structural Requirements for Activity of Mind bomb1 in Notch Signaling"

### Supplementary Fig. 1

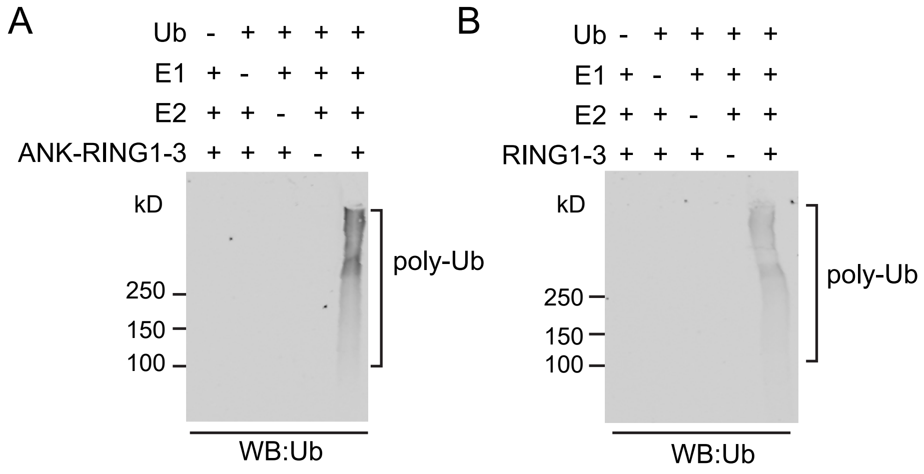

### Supplementary Fig. 2

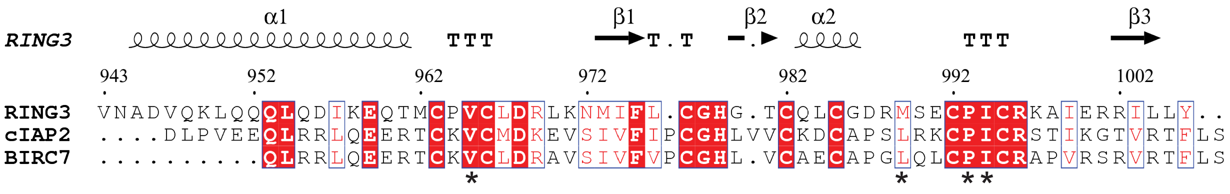

Supplementary Fig. 3

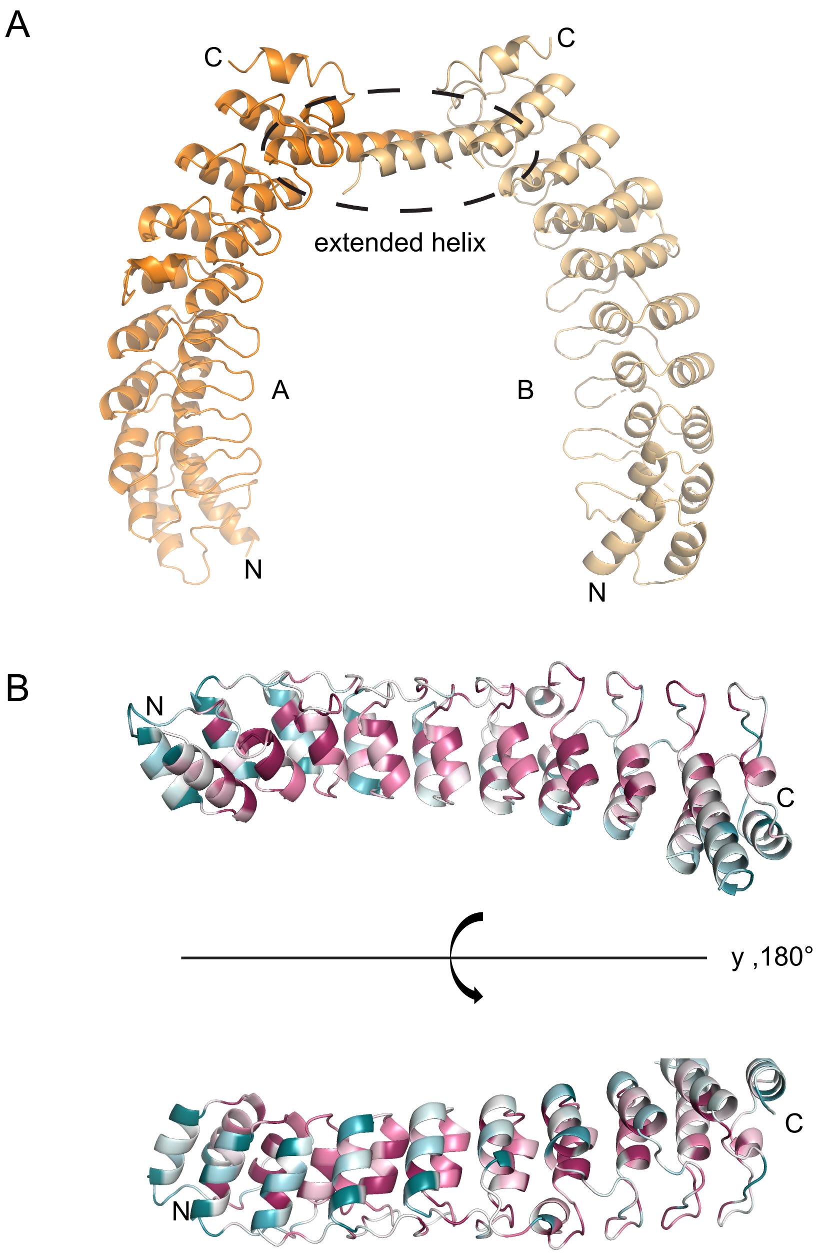

Supplementary Fig. 4

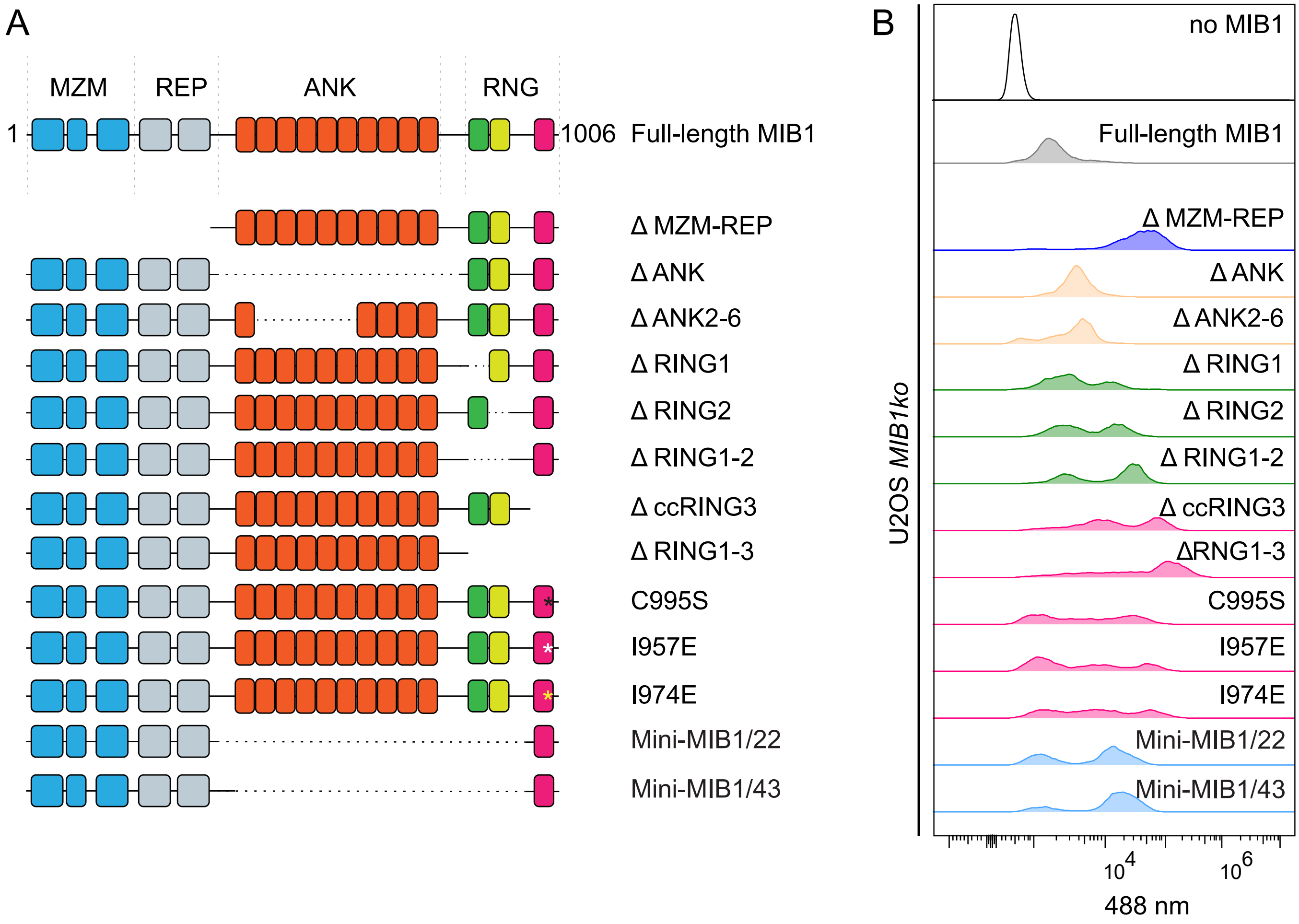

Supplementary Fig. 5

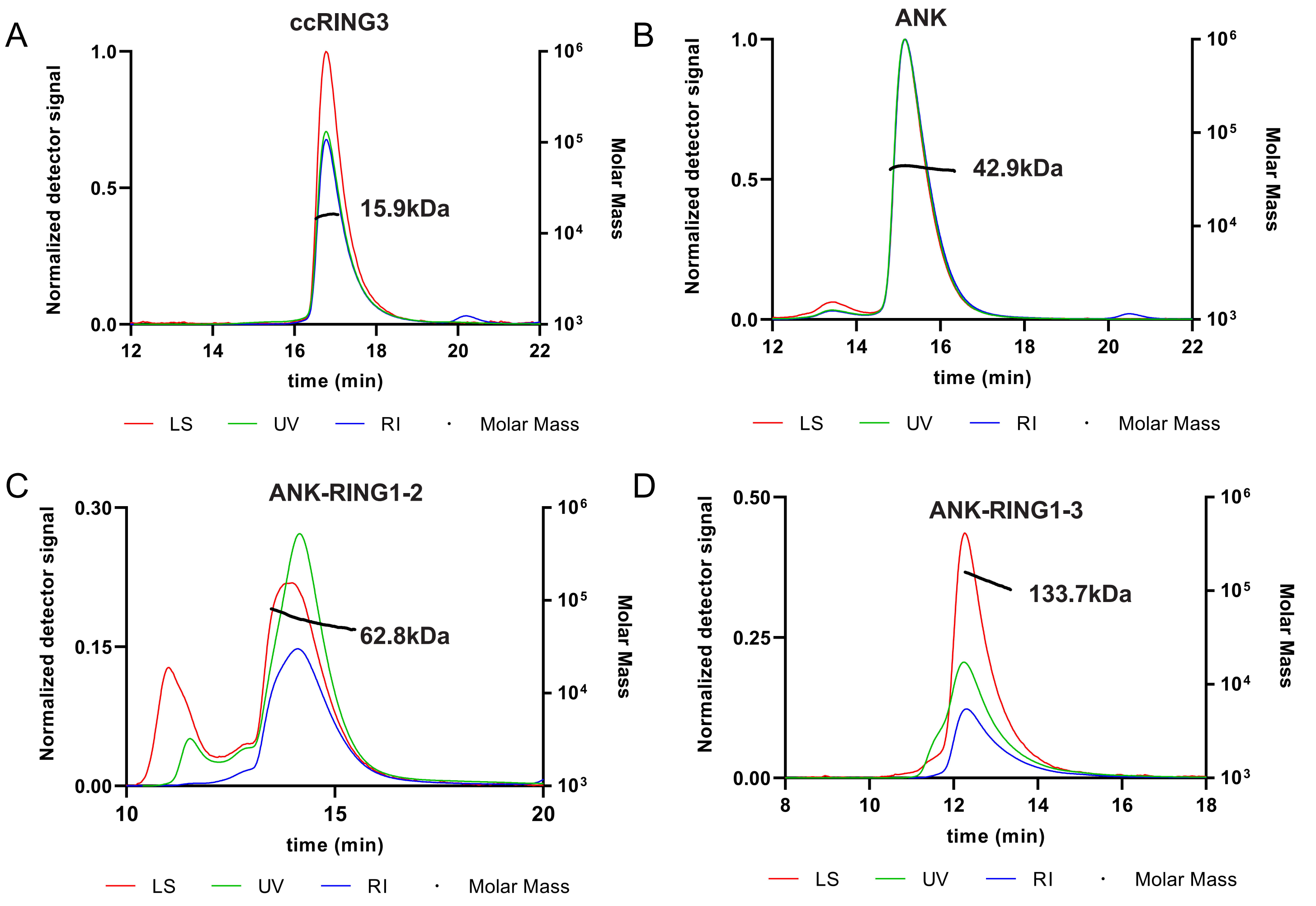

Supplementary Fig. 6

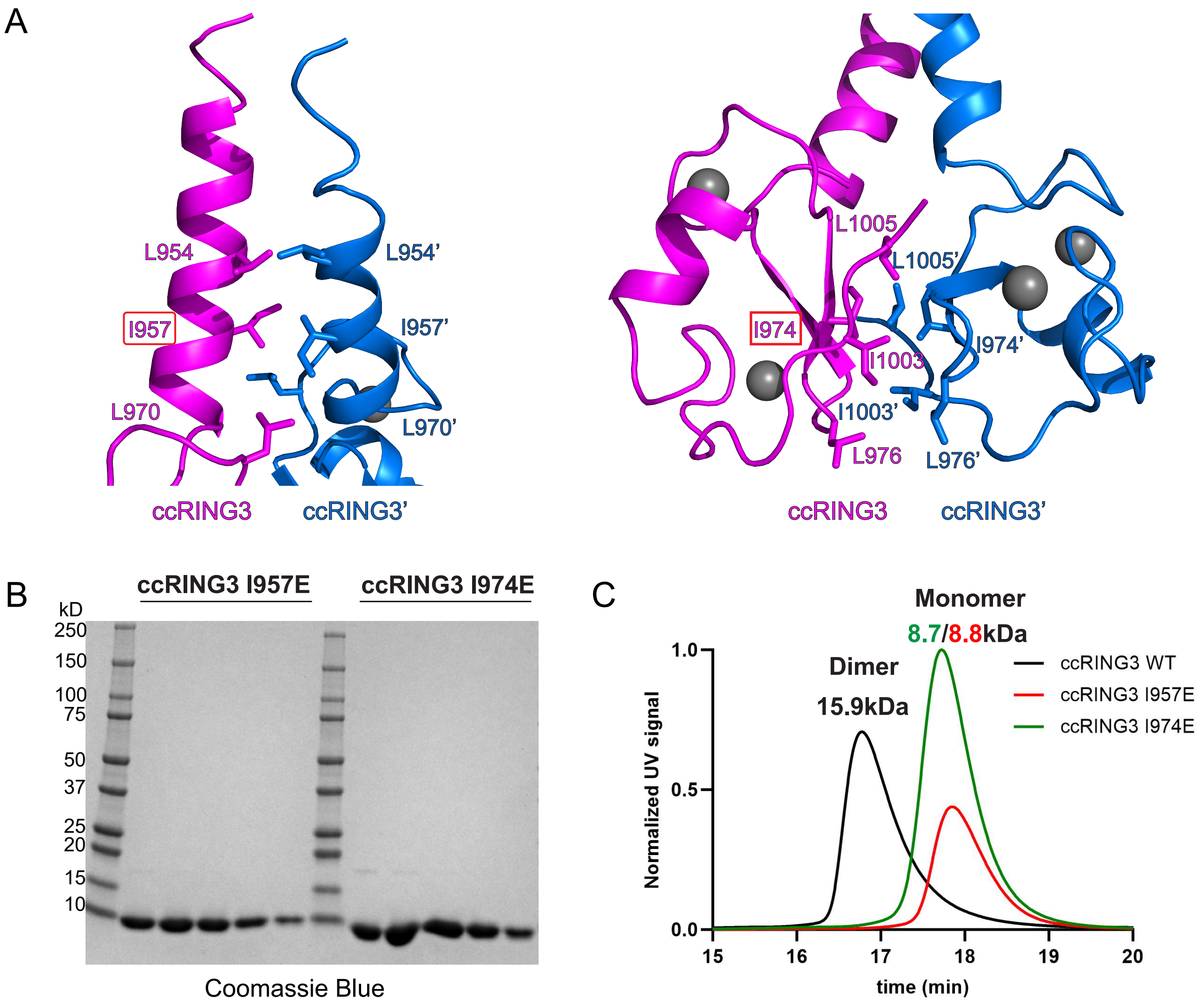
